# Brain Pretargeted PET – New Horizons to Image CNS Targets with Monoclonal Antibodies

**DOI:** 10.1101/2024.10.12.617694

**Authors:** Tobias Gustavsson, Thomas Kustermann, Lars Hvass, Vladimir Shalgunov, Anne Skovsbo Clausen, Sophie Stotz, Thomas Erik Wünsche, Gitte M. Knudsen, Blanca I. Aldana, Jens Niewoehner, Luca Gobbi, Michael Honer, Umberto Battisti, Andreas Kjaer, Matthias M. Herth

## Abstract

Antibodies are excellent targeting vectors for molecular imaging. Slow pharmacokinetics and low blood-brain barrier penetration hinder their widespread application for molecular imaging within CNS. Improved brain uptake can be achieved via transferrin-mediated transcytosis. Pretargeted imaging can increase imaging contrast and reduce radiation exposure to the patient. Here, we report for the first time that pretargeted imaging of CNS targets using intravenously administered target vectors is feasible. Specific binding to Aβ-bound antibody was achieved. We believe that this proof-of-concept study will facilitate molecular imaging of currently ‘undruggable’ targets with antibodies where small molecule PET tracer discovery has been challenging.

## Introduction

Drug development for many neurodegenerative diseases such as Alzheimer’s disease or Parkinson’s disease has proven complex over the past decades. A major obstacle to progress is the lack of suitable biomarkers. For many therapeutic targets, it remains challenging to demonstrate target engagement or impact on disease progression, partly due to the lack of suitable PET tracers. Amyloid beta (Aβ), tau or α-synuclein in its various protofibril and (iso)forms remain some of the major therapeutic and biomarker targets for proteinopathies. While there are validated PET tracers for proteins such as aggregated Aβ and 3R/4R tau, the sensitivity for structural differences in pathologies species (e.g. oligomers, protofibrils and fibrils) remains unknown. For other targets, such as aggregated forms of α-synuclein or TDP-43, no suitable PET tracers exist. Increasing focus on other pathogenic factors like neuroinflammation, mitochondrial dysfunction, or synaptic dysfunction has highlighted the need to develop tracers for other proteins, particularly for those considered ‘undruggable’, i.e. for proteins where small molecule PET tracer discovery has been challenging. High molecular weight binders such as monoclonal antibodies (mAbs) with their excellent target specificity along with improved brain penetration upon modification, e.g. by conjugating a transferrin receptor targeting moiety, may offer an alternative imaging approach.^1-4^ This approach bears the promise for repurposing existing libraries of antibody constructs for PET imaging, thereby considerably reducing the need for de novo PET tracer discovery programs. Recent preclinical studies have shown that antibody-based PET imaging is feasible for targets beyond the blood-brain barrier (BBB).^1-3^ However, the long biological half-life of antibodies requires PET radionuclides with long decay half-lives. The use of such radionuclides poses challenges to obtain good signal-to-background ratios in PET measurements. Typically, these radionuclides also result in high radiation burden to patients.^3-7^ Shorter lived and clinically used PET radionuclides such as fluorine-18 are unsuitable to directly image target binding of mAbs.^4^ In contrast, pretargeted imaging, based on bioorthogonal click chemistry between a target-bound antibody and a small molecule radiotracer, offers the possibility to use short-lived radionuclides to image target binding of mAbs, increase signal-to-background ratios and reduce radiation exposure.^5, 6^

Compared to conventional antibody PET imaging where antibodies are directly labeled with long-lived radionuclides, pretargeting allows PET imaging with antibodies using short-lived radionuclides.^6^ Pretargeting uncouples the typically long biological half-life of antibodies from the radionuclide decay half-life, i.e. short-lived radionuclides can be used to image slow accumulating and excreting mAbs. Pretargeting combines the exquisite affinity and specificity of mAbs with the optimal pharmacokinetics of small molecules resulting in high imaging contrast and low radiation exposure to healthy tissue.^6^ Several pretargeting strategies are based on bioorthogonal reactions.^7,8^ The most prominent bioorthogonal reaction is the tetrazine ligation between a trans-cyclooctene (TCO) and a tetrazine (Tz). This ligation is highly selective and ultrafast (up to 10^7-9^ M^-1^s^-1^), irreversible and results in a stable product.^6-11^

While several examples of pretargeted tumor imaging have been reported,^10,14^ successful application of pretargeted imaging in the brain remains elusive, except for instances where pretargeted vectors have been directly administered via intrathecal or intracerebral routes.^12-17^ It would be highly beneficial for drug development and therapy monitoring to have an imaging method available to quantify BBB penetrant mAbs intravenously administered. Multiple antibodies are currently under development to engage ‘undruggable’ brain targets with the aim to develop next-generation therapies. In 2023, Roche reported the development of a bispecific antibody with a Brain Shuttle™ module.^15^ This module allows for facilitated transport of mAbs across the BBB using transferrin-mediated transcytosis. In the same year, the Herth group developed a ^18^F-labeled Tz that could successfully be used in vivo to image an intracerebral deposited TCO-polymer.^13^ The Tz displayed optimal properties for pretargeted brain imaging. The in vivo clicked Tz showed high brain uptake and accumulation as well as fast blood and brain clearance of unclicked Tz.^13^ In the present proof-of-concept study, we report for the first time that pretargeted imaging of an intravenously injected TCO-tagged antibody is possible in vivo by combining the antibody brain shuttle module with a BBB penetrant ^18^F-labeled Tz (**Figure 1**).

**Figure 1.**
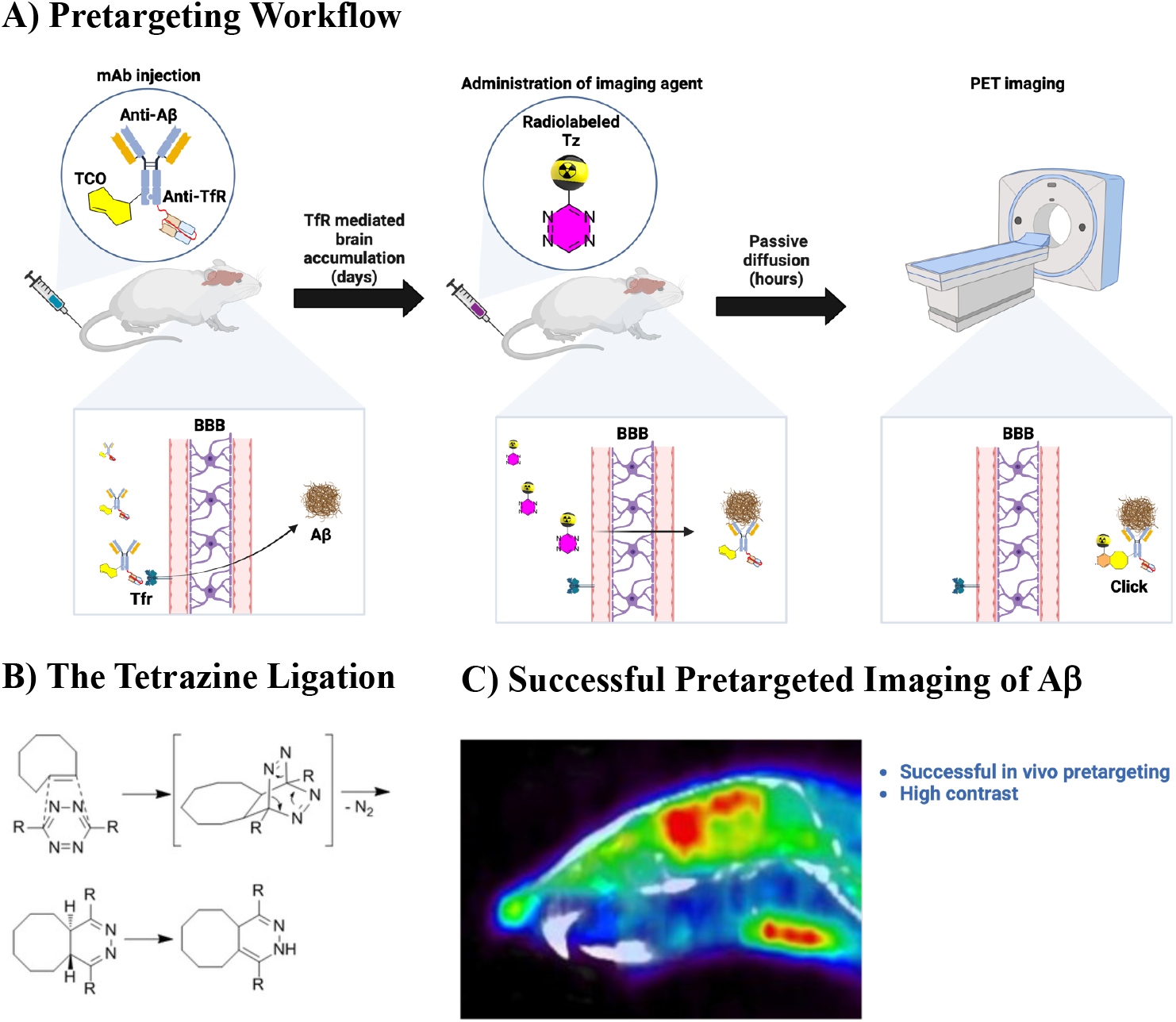
**A)** Workflow of pretargeted PET imaging beyond the blood-brain barrier (BBB), **B)** Tetrazine (Tz) ligation, **C)** Amyloid beta (Aβ) pathology was successfully visualized by a pretargeting approach. PET was conducted using the pretargeting vector, TCO-BS-mAb31, and the imaging agent, [^18^F]**1**. TCO-BS-mAb31 was administered 72h before the imaging agent.

### Chemistry and mAb characterization

As a model antibody, we used the Aβ binding antibody, mAb31, in its brain shuttle version (BS-mAb31), modified it with TCOs and imaged it with [^18^F]**1**. The bispecific antibody construct, BS-mAb31, targeting fibrillar Aβ and the murine transferrin receptor (mTfR), was supplied by Roche.^15^ BS-mAb31 was TCO-functionalized with NHS-TCO (2.2 mg BS-mAb31 (11 nmol) was incubated with a 20:1 mole ratio of NHS-TCO:BS-mAb31 in PBS (pH 8.4, 650 μL)). TCO quantification according to Shulganov et al.^13^ revealed an active TCO-load of 9.5 TCOs per BS-mAb. ELISA measurements showed that this modification did not alter the binding affinity of TCO-modified BS-mAb31(TCO-BS-mAb31) towards mTfR and protofibrillar Aβ compared to non-modified BS-mAb. No aggregation of TCO-BS-mAb31 was observed via HPLC-SEC. [^18^F]**1** was radiolabeled as reported by Shulganov et al.^13^

### Pretargeted PET imaging

Evaluation studies were performed in 5 months old female 5xFAD and wildtype (WT) mice. Three days prior to administration of [^18^F]**1**, 5xFAD mice were intravenously injected with TCO-BS-mAb at a dose of 41.4 ± 2.7 pmol/g (mean ± SD). The corresponding dose for WT mice was 38.2 ± 1.1 pmol/g. TCO-ligation of [^18^F]**1** within the brain were evaluated during a 90 min dynamic PET scan. 5xFAD mice received 8.7 ± 1.6 MBq of [^18^F]**1**, while WT mice received 8.6 ± 1.1 MBq.

PET data clearly demonstrated that 5xFAD mice had a high retention of [^18^F]**1** in Aβ-rich areas, including the cortex and hippocampus (in contrast to WT mice) (**Figure 2A**). Quantified time activity curves (TACs) demonstrated that 5xFAD mice had 30-50% higher [^18^F]**1** retention in selected Aβ-rich regions then WT mice (**Figure 2B**). 5xFAD mice exhibited a low Aβ burden in the cerebellum, similar to the uptake of [^18^F]**1** in WT mice. These results indicate the suitability of the cerebellum as a reference region for PET data quantification. Overall, increases in the regions-to-cerebellum ratio were particularly pronounced in 5xFAD mice (**Figure 2C**), demonstrating that administration of TCO-BS-mAb31 and subsequent in vivo labeling by [^18^F]**1** allows specific visualization of brain Aβ pathology.

**Figure 2.**
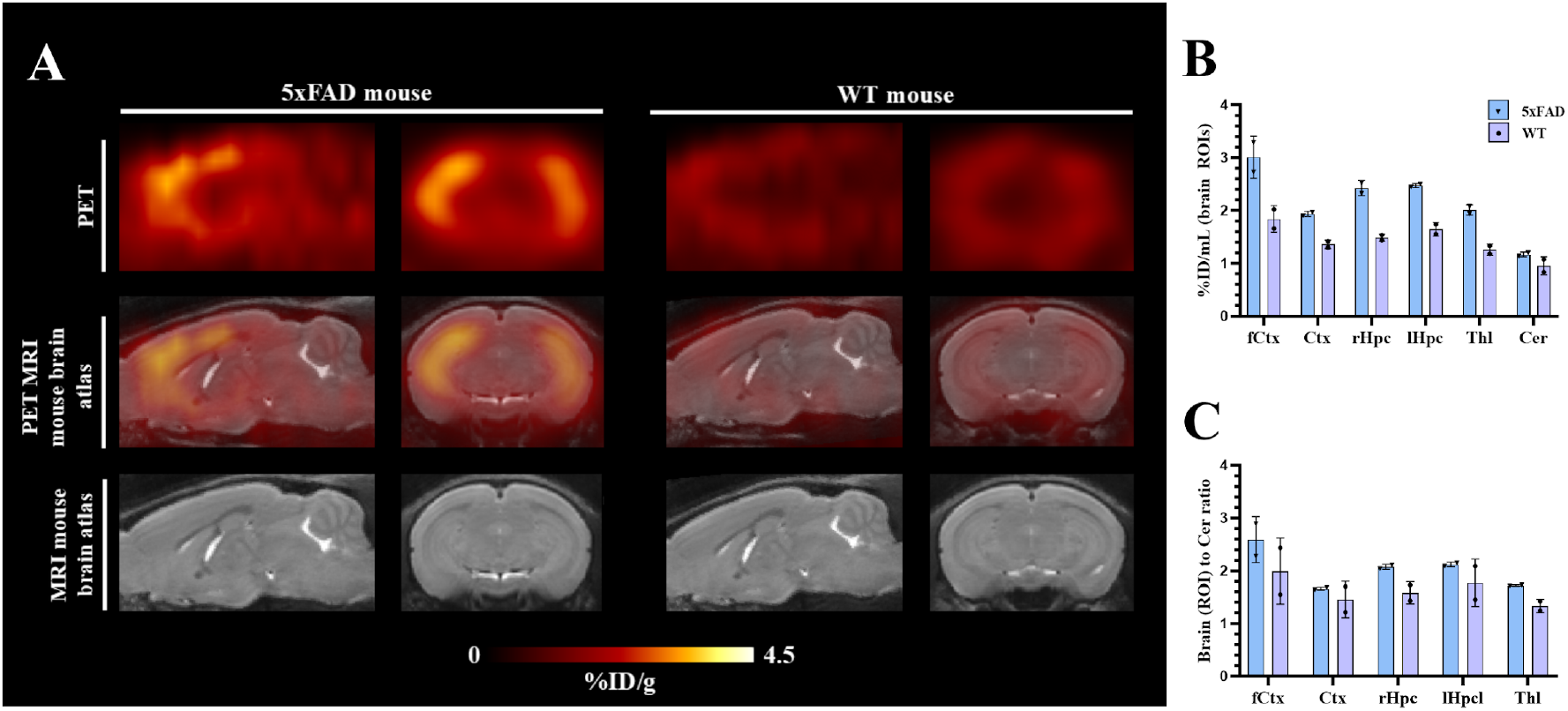
**A**) Dynamic PET data (80-90 min) showing sagittal and coronal brain projections of selected 5xFAD and WT mice overlayed with mouse brain MRI template from Presotto et al.^18^ **B**) Quantified TAC (80-90 min) extracted from dynamic PET using volume-of-interest (VOIs) from Mirrione et al.^19^ and expressed as %ID/mL. **C**) Regions (ROI)-to-cerebellum ratio. Values are shown as mean and error bars are standard deviation. fCtx, frontal cortex; Ctx, cortex; rHpc, right hippocampus; lHpc, left hippocampus; Thl, thalamus; Cer, cerebellum.

## Conclusion

Here, we report for the first time that pretargeted imaging of CNS targets using intravenously administered target vectors is feasible. Specific binding to Aβ-bound antibody was achieved. Pretargeted imaging has the potential to unlock CNS PET imaging for a wide variety of previously intractable targets by taking advantage of the high sensitivity and specificity of immune-based targeting vectors. Antibodies already developed for disease-relevant targets could be repurposed for pretargeted PET imaging, thereby accelerating the development of CNS PET diagnostics.

